# Comparison of test-retest reproducibility of DESPOT and 3D-QALAS for water *T*_1_ and *T*_2_ mapping

**DOI:** 10.1101/2024.08.15.608081

**Authors:** Gizeaddis Lamesgin Simegn, Borjan Gagoski, Yulu Song, Douglas C. Dean, Kathleen E. Hupfeld, Saipavitra Murali-Manohar, Christopher W. Davies-Jenkins, Dunja Simičić, Jessica Wisnowski, Vivek Yedavalli, Aaron T. Gudmundson, Helge J. Zöllner, Georg Oeltzschner, Richard A. E. Edden

## Abstract

**Purpose:** Relaxometry, specifically *T*_1_ and *T*_2_ mapping, has become an essential technique for assessing the properties of biological tissues related to various physiological and pathological conditions. Many techniques are being used to estimate *T*_1_ and *T*_2_ relaxation times, ranging from the traditional inversion or saturation recovery and spin-echo sequences to more advanced methods. Choosing the appropriate method for a specific application is critical since the precision and accuracy of *T*_1_ and *T*_2_ measurements are influenced by a variety of factors including the pulse sequence and its parameters, the inherent properties of the tissue being examined, the MRI hardware, and the image reconstruction. The aim of this study is to evaluate and compare the test-retest reproducibility of two advanced MRI relaxometry techniques (Driven Equilibrium Single Pulse Observation of *T*_1_ and *T*_2_, DESPOT, and 3D Quantification using an interleaved Look-Locker acquisition Sequence with a *T*_2_ preparation pulse, QALAS), for *T*_1_ and *T*_2_ mapping in a healthy volunteer cohort.

**Methods:** 10 healthy volunteers underwent brain MRI at 1.3 mm^3^ isotropic resolution, acquiring DESPOT and QALAS data (∼11.8 and ∼5 minutes duration, including field maps, respectively), test-retest with subject repositioning, on a 3.0 Tesla Philips Ingenia Elition scanner. To reconstruct the *T*_1_ and *T*_2_ maps, we used an equation-based algorithm for DESPOT and a dictionary-based algorithm that incorporates inversion efficiency and *B_1_*-field inhomogeneity for QALAS. The test-retest reproducibility was assessed using the coefficient of variation (CoV), intraclass correlation coefficient (ICC) and Bland-Altman plots.

**Results:** Our results indicate that both the DESPOT and QALAS techniques demonstrate good levels of test-retest reproducibility for *T*_1_ and *T*_2_ mapping across the brain. Higher whole-brain voxel-to-voxel ICCs are observed in QALAS for *T*_1_ (0.84 ± 0.039) and in DESPOT for *T*_2_ (0.897 ± 0.029). The Bland-Altman plots show smaller bias and variability of *T*_1_ estimates for QALAS (mean of -0.02 s, and upper and lower limits of -0.14 and 0.11 s, 95% CI) than for DESPOT (mean of -0.02 s, and limits of -0.31 and 0.27 s). QALAS also showed less variability (mean 1.08 ms, limits –1.88 to 4.04 ms) for *T*_2_ compared to DESPOT (mean of 2.56 ms, and limits -17.29 to 22.41 ms). The within-subject CoVs for QALAS range from 0.6% (*T*_2_ in CSF) to 5.8% (*T*_2_ in GM), while for DESPOT they range from 2.1% (*T*_2_ in CSF) to 6.7% (*T*_2_ in GM). The between-subject CoVs for QALAS range from 2.5% (*T*_2_ in GM) to 12% (*T*_2_ in CSF), and for DESPOT they range from 3.7% (*T*_2_ in WM) to 9.3% (*T*_2_ in CSF).

**Conclusion:** Overall, QALAS demonstrated better reproducibility for *T*_1_ and *T*_2_ measurements than DESPOT, in addition to reduced acquisition time.

## 1. Introduction

MRI relaxation times are crucial parameters for tissue characterization. The longitudinal relaxation time, or *T*_1_, characterizes the exponential restoration of equilibrium spin-state populations. The transverse, or *T*_2_, relaxation time characterizes the exponential loss of phase coherence. *T*_1_ and *T*_2_ index different tissue properties and can be mapped across the brain using quantitative experiments that sample different stages of the relaxation processes. Quantitative assessment of *T*_1_ and *T*_2_ can enhance the diagnostic accuracy beyond conventional relaxation-weighted MRI in applications ranging from the detection and assessment of myocarditis (1–4), liver fibrosis and cirrhosis (5–9), cartilage degeneration to monitoring disease progression and response to therapy (10–12), neurologic disease (13–17), normal brain development (18,19), and aging (20–22). Quantitative MRI also allows for the objective comparison of pathological conditions across time and between individuals, making it an indispensable tool in both research and clinical settings.

The simplest, and gold standard, method for generating *T*_1_ maps involves acquiring images at multiple inversion times (TI), i.e. inversion recovery (IR) sequences, and modeling voxel-by-voxel with a *T*_1_ decay curve. However, IR sequences require very long repetition times (TR), resulting in scan times that are impractical for clinical use and may also lead to inaccuracies due to patient motion (23). The gold standard for generating *T*_2_ maps involves multi-echo spin-echo sequences. Several techniques for more rapid *T*_1_ mapping and combined *T*_1_-*T*_2_ mapping have been developed, including the modified look-locker IR (MOLLI) (24), shortened MOLLI (shMOLLI) (25), saturation recovery single-shot acquisition (SASHA) (26), True *T*_1_ mapping with SMAR*T*_1_Map (27), TrueFISP (28), variable flip angle imaging (29,30), magnetization-prepared 2 rapid gradient echo (MP2RAGE) (31), and multi-echo (ME) extension of MP2RAGE (MP2RAGEME) (32). Most of these techniques are either designed for *T*_1_ mapping only, require a long acquisition time or are 2D mapping methods, which limits their utility in. Clinical and research settings.

Driven Equilibrium Single Pulse Observation of *T*_1_ and *T*_2_ (DESPOT) (33,34) and 3D quantification using an interleaved Look-Locker acquisition sequence with a *T*_2_ preparation pulse (QALAS) (35) are two relatively new, fast 3D techniques that allow simultaneous mapping of *T*_1_ and *T*_2_ at high spatial resolution. DESPOT includes DESPO*T*_1_ and DESPO*T*_2_ for *T*_1_ and *T*_2_ mapping, respectively. DESPO*T*_1_ acquires a series of spoiled gradient recalled-echo (SPGR) images with the same TR and different flip angles, while DESPO*T*_2_ acquires fully balanced steady-state free precession (bSSFP) images at different flip angles with constant TR (33,34). QALAS, a newer technique, is based on 3D spoiled Turbo Field Echo sequences using inversion recovery interleaved *T*_2_ preparation. The QALAS acquisition consists of five turbo-FLASH readouts. A *T*_2_-preparation module precedes the first readout, followed by an inversion pulse, so that the following four readouts capture *T*_1_ dynamics (35). It combines *T*_1_, *T*_2_, and proton density (PD) mapping and evaluation of inversion efficiency (IE) in a single acquisition.

Measurements of *T*_1_ and *T*_2_ maps can be influenced by various factors, including the sequence type, acquisition sampling scheme, reconstruction technique, magnetization transfer, flow effects, *T*_2_ effects, and motion (23,36–38). Even for “gold standard” methods, measurements depend on the choice of inversion or echo times sampled. Establishing the reproducibility and accuracy of *T*_1_ and *T*_2_ mapping is crucial for their adoption in clinical practice and longitudinal studies, particularly for newer ‘hybrid’ methods, like DESPOT and QALAS, that map *T*_1_ and *T*_2_ from a complex set of mixed-contrast images. The purpose of this study is to evaluate and compare the test-retest reproducibility of *T*_1_ and *T*_2_ mapping using DESPOT and QALAS techniques.

## 2. Methods

### 2.1. MRI Acquisition

Data were acquired from 10 healthy volunteers (5 male, 5 female, ages 23 to 49) after obtaining informed written consent. The study was approved by the Johns Hopkins University Institutional Review Board. All scans were conducted using a 3.0 Tesla Philips Ingenia Elition MRI scanner, equipped with a 32-channel receive head coil. For anatomical co-registration, we first collected a *T*_1_-weighted structural MPRAGE scan using the following parameters: TR / TE 2000 ms / 2 ms, flip angle 8°, 150 1-mm slices, voxel size 1 mm^3^ isotropic, total time 2 min 46 sec. Then, we acquired DESPOT images at 1.3 mm^3^ isotropic resolution (FOV 224×22×x166 mm^3^) using an SPGR sequence with flip angles of 4°, 12°, and 18° and TR / TE of 6.3 ms / 3 09 ms, and a bSSFP sequence with flip angles of 15°, 30°, and 60° TR / TE of 6.3 ms / 3.09ms . bSSFP images were acquired with phase cycling patters of 0° and 180° to allow correction for main magnetic field (*B*_0_) inhomogeneities (39). A *B*_1_ map (40) was also acquired. The total acquisition time for DESPOT including the filed map was 11.87 minutes. Similarly, QALAS was acquired with TR / TE of 5.7 ms / 2.3 ms, 1.3 mm^3^ isotropic resolution, FOV 224×224×166 mm^3^, flip angle 4°. The five turbo-FLASH readouts were spaced 900 ms apart, and a 100-ms *T*_2_-preparation module was used. Our QALAS protocol (which, of note, is the same as that used in the HEALthy Brain and Child Development (HBCD) (41) study actively collecting longitudinal neuroimaging data from 7,500 children) also acquires *B*_1_ maps, using the 48.4 s long actual flip-angle imaging (AFI (40)) sequence. The total acquisition time for QALAS including the field map was 5.03 minutes.

### 2.2. *T*_1_ and *T*_2_ reconstruction and mapping

#### 2.2.1. DESPOT

Following acquisition, SPGR, bSSFP and *B*_1_ map, were linearly co-registered to account for subtle head movement (42) and non-parenchyma signal was removed (39). The rapid combined *T*_1_ and *T*_2_ mapping approaches called DESPO*T*_1_ and DESPO*T*_2_-FM, respectively were used (33). *T*_1_ is estimated from the SPGR data acquired at different flip angles, modeled according to:

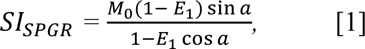

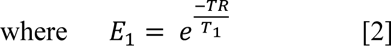

where *SI_SPGR_* is the SPGR signal intensity as a function of flip angle *a*, *M*_2_ is a factor proportional to the equilibrium longitudinal magnetization, and *E*_’_ is expressed in Equation 2. Similarly, *T*_2_ is estimated by fitting the bSSFP images acquired at three different flip angles to the following:

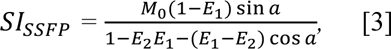

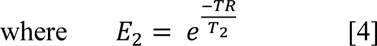

*SI*_SSFP_ is the SSFP signal intensity associated with flip angle α. NIFTI-format *T*_1_ and *T*_2_ maps were generated in a python environment using the qmri-neuropipe tool (43).

#### 2.2.2. QALAS

To estimate the QALAS *T*_1_ and *T*_2_ maps, a dictionary-based matching algorithm (44,45) that incorporates Inversion Efficiency (IE) and *B_1_* field corrections (46) was used. The dictionary encompassed IE values ranging from 0.75 to 1, in 10 equal steps, *B_1_* field corrections ranging from 0.65 to 1.35, in 25 equal steps, *T*_1_ ranges (5:2:1200 ms, 1200:5:2100 ms, 2100:10:3100 ms, 3100:20:5000 ms), and *T*_2_ ranges (1:2:150 ms, 150:5:360 ms, 3760:10:1100 ms, 1100:50:2500 ms). The dictionary was generated from simulations of the five sub-experiments using MATLAB R2023a (The MathWorks, Natick, MA) on a high-capacity server cluster. The signals after the inversion recovery were adjusted based on their respective IE values. The *B_1_*^+^ inhomogeneity maps generated were applied to the flip angles within the turbo-flash readouts to account for spatial variations. The dictionary-matching used to generate *T*_1_ and *T*_2_ maps in NIfTI format was also performed using MATLAB R2019b on the same server clusters.

### 2.3. Postprocessing

All DESPOT and QALAS test and retest T1 and T2 maps were first co-registered and resliced to the high-resolution *T*_1_-weighted image (MP-RAGE). GM, WM and CSF masks were derived from SPM 12 segmentation (47) of the *T*_1_-weighted MP-RAGE (with a probability threshold of 0.8) and subsequently applied to the *T*_1_ and *T*_2_ maps. All post-processing, including image co-registration, re-slicing, segmentation, and brain region masking was performed using MATLAB R2023a. Figure 1 shows the post-processing procedures.

**Figure 1:**
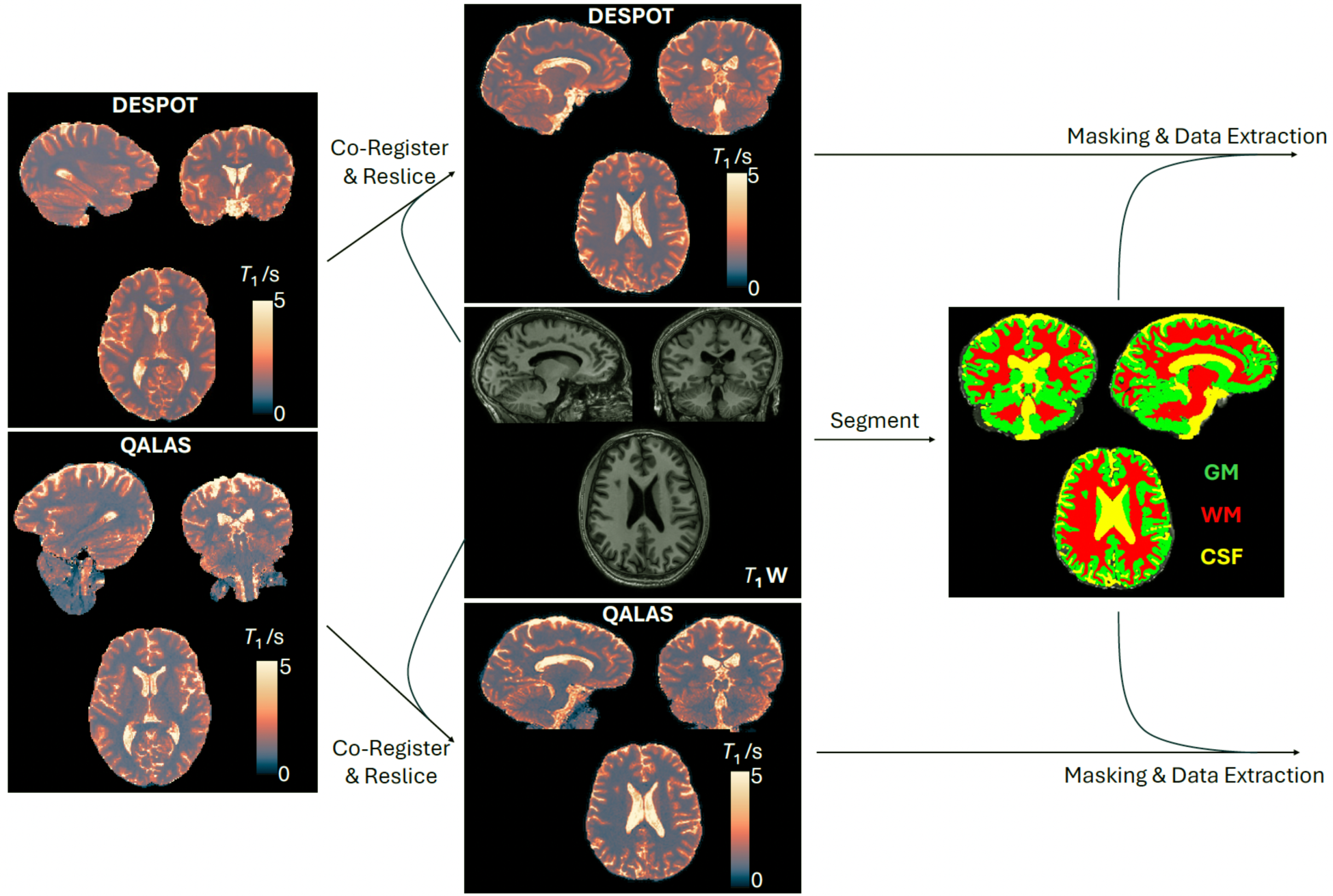
Post-processing procedure. Both the DESPOT and QALAS test-retest T_1_ and T_2_ maps were co-registered and resliced to the same reference T_1_-weighted image (T_1_w). The T_1_-weighted image was segmented (which includes automatic skull stripping), each segmented regions (GM, WM and CSF) were binarized after 90% thresholding using SPM12 tool to generate the masks. The masks were then applied to the co-registered and resliced maps for data extraction.

### 2.4. Statistical Analysis

After segmentation and masking, histograms were plotted for *T*_1_ and *T*_2_ measurements across the GM, WM and CSF masks. The average *T*_1_ and *T*_2_ within each tissue mask was plotted for the test and retest acquisitions of each subject. These values are then used to generate within-subject and between-subject coefficients of variation (CoVs) for test-retest acquisitions, according to:

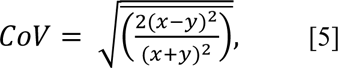

where the bar represents a mean across subjects, and x and y are the test and retest values, respectively. The between-subject CoV is determined from the ratio of standard deviations of test-retest subject mean values and average of these test-retest subject mean values. The test-retest reproducibility of DESPOT and QALAS was separately visualized with Bland-Altman plots of the tissue-mask values (x and y) values. Voxel-by-voxel reproducibility of the maps was examined using ICCs to assess the reliability and reproducibility of *T*_1_ and *T*_2_ measurements across the test-retest sessions.

In addition to test and retest evaluation of each method, the agreement between DESPOT and QALAS measurements was visualized using Bland-Altman plots, plotting the difference between DESPOT and QALAS ‘test’ measurements against the average of the two ‘test’ measurements (i.e. ignoring retest for both methods). Voxel-to voxel ICCs were also evaluated between the two methods for ‘test’ measurements. This analysis provided insights into the bias and limits of agreement between the techniques.

## 3. Results

### 3.1. Regional *T*_1_ and *T*_2_ values

Histograms of *T*_1_ and *T*_2_ derived from DESPOT and QALAS for a representative subject, for each segmented tissue type, are shown in Figure 2. Test and retest generally show excellent agreement for each of the two methods. While there is slight disagreement between DESPOT and QALAS for *T*_1_ and *T*_2_ of GM and WM tissues, this is much more pronounced for CSF. The mean *T*_1_ and *T*_2_ values across GM, WM, and CSF masks for test-retest measurements using DESPOT and QALAS is shown for all 10 subjects in Figure 3.

**Figure 2:**
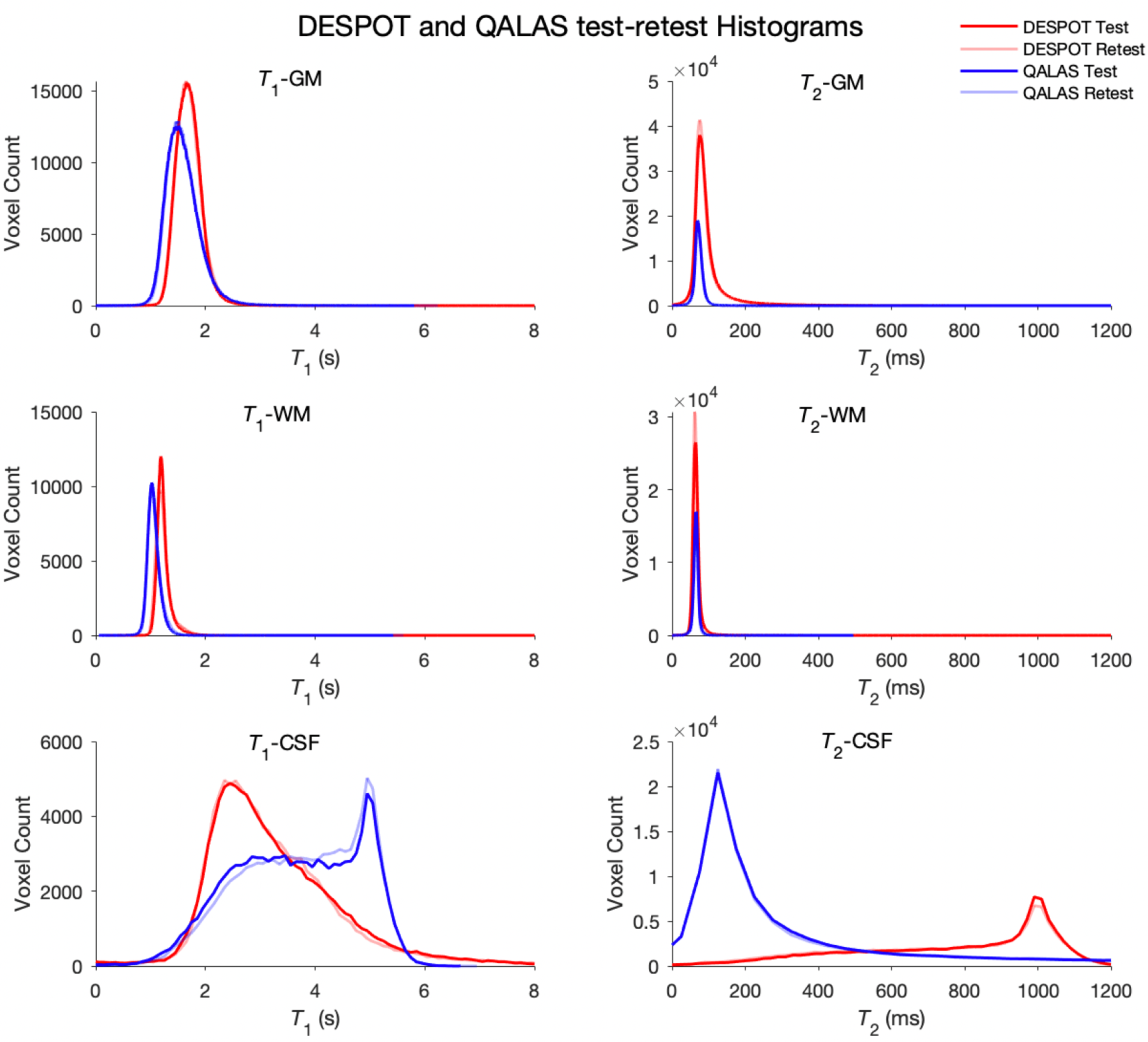
Single subject DESPOT and QALAS test-retest T_1_ and T_2_ maps overlaid histogram plots for GM, WM, and CSF.

**Figure 3:**
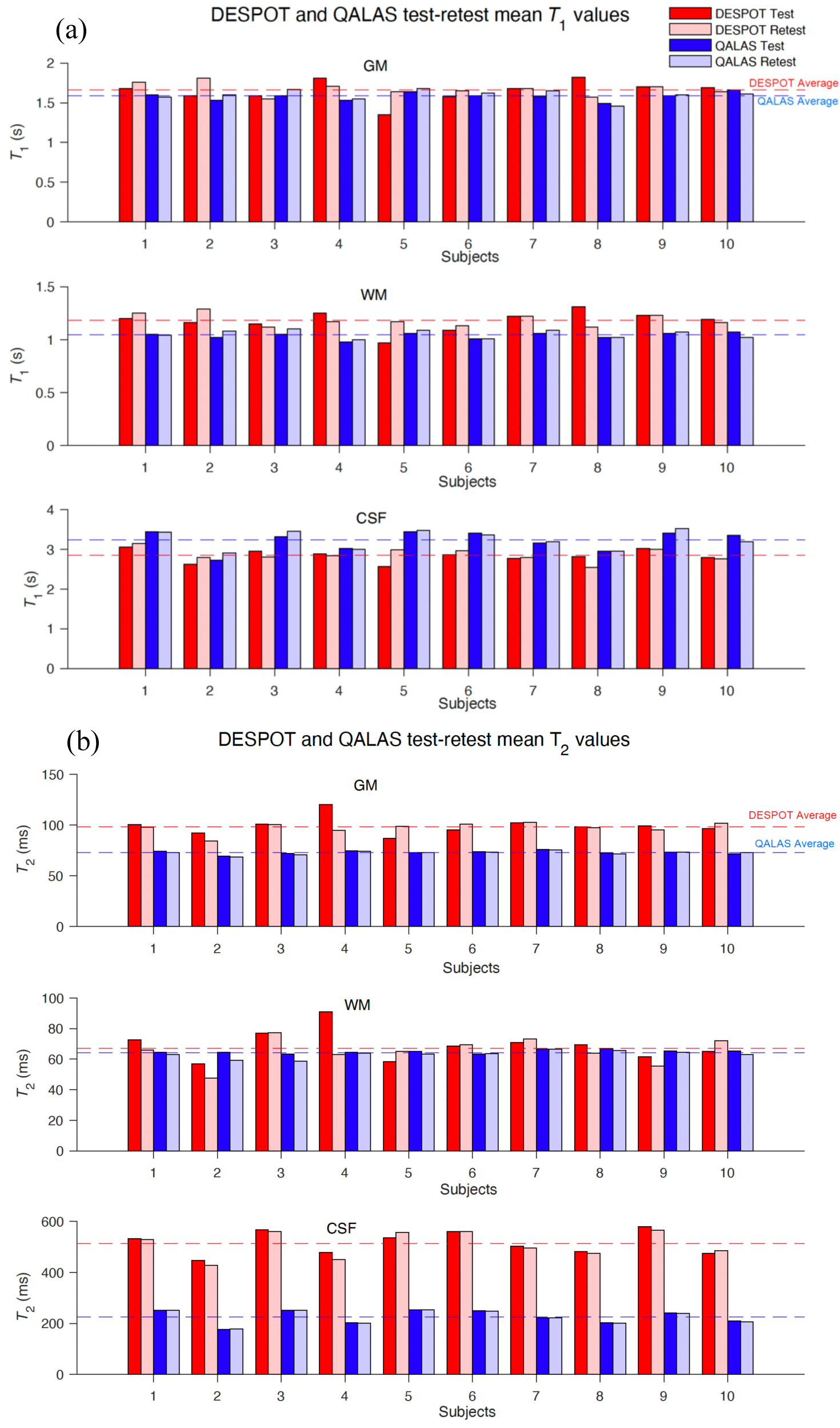
DESPOT and QALAS test-retest GM, WM and CSF mean of (a) T_1_ and (b) T_2_ values.

#### 3.2. Coefficient of Variations (CoVs)

Table 1 shows the within- and between-subject CoVs for mean *T*_1_ and *T*_2_ values across tissue masks, for DESPOT and QALAS. For DESPOT, the within-subject CoVs vary from a minimum of 2.1% for *T*_2_ in CSF to a maximum of 6.7% for *T*_2_ in GM. For QALAS, CoVs vary from a minimum of 0.6% for *T*_2_ in CSF to a maximum of 5.8% for *T*_2_ in WM. The between-subject CoVs for DESPOT range from 3.7% for *T*_2_ in WM, to 9.3% for *T*_2_ in CSF, and for QALAS range from a minimum of 2.5% for *T*_2_ in GM to a maximum of 12% for *T*_2_ in CSF. In almost all cases, CoVs were smaller for QALAS than DESPOT.

**Table 1:** CoVs for mean T_1_ and T_2_ Values in GM, WM, and CSF using DESPOT (D) and QALAS (Q) in Test-Retest Measurements.

| CoV (%) | $T_1$ | | | | | | $T_2$ | | | | | |
| --- | --- | --- | --- | --- | --- | --- | --- | --- | --- | --- | --- | --- |
|  | GM |  | WM |  | CSF |  | GM |  | WM |  | CSF |  |
|  | D | Q | D | Q | D | Q | D | Q | D | Q | D | Q |
| Within-subject | 6.5 | 2.1 | 6.3 | 2.2 | 4.6 | 1.9 | 6.7 | 2.2 | 5.1 | 5.8 | 2.1 | 0.6 |
| Between-subject | 4.7 | 3.3 | 4.8 | 2.8 | 4.6 | 7.3 | 5.2 | 2.5 | 3.7 | 2.6 | 9.3 | 12.0 |

#### 3.3. Test-Retest Bland-Altman plots

DESPOT and QALAS Bland-Altman plots for GM, WM and CSF *T*_1_ and *T*_2_ values of 10 subjects are shown in Figure 4. The DESPOT *T*_1_ plot shows a mean bias of –0.02 s and relatively wide 95% confidence intervals (–0.31 to 0.27 s). The QALAS *T*_1_ plot shows a similar bias of –0.02 s with narrower 95% confidence intervals (–0.14 to 0.11 s), indicating good consistency. For *T*_2_ measurements, the DESPOT plot shows a mean difference of 2.56 ms with a limit of agreement from –17.29 to 22.41 ms, while the QALAS plot exhibits a smaller mean difference of 1.08 ms and narrower limits of agreement (–1.88 to 4.04 ms).

**Figure 4:**
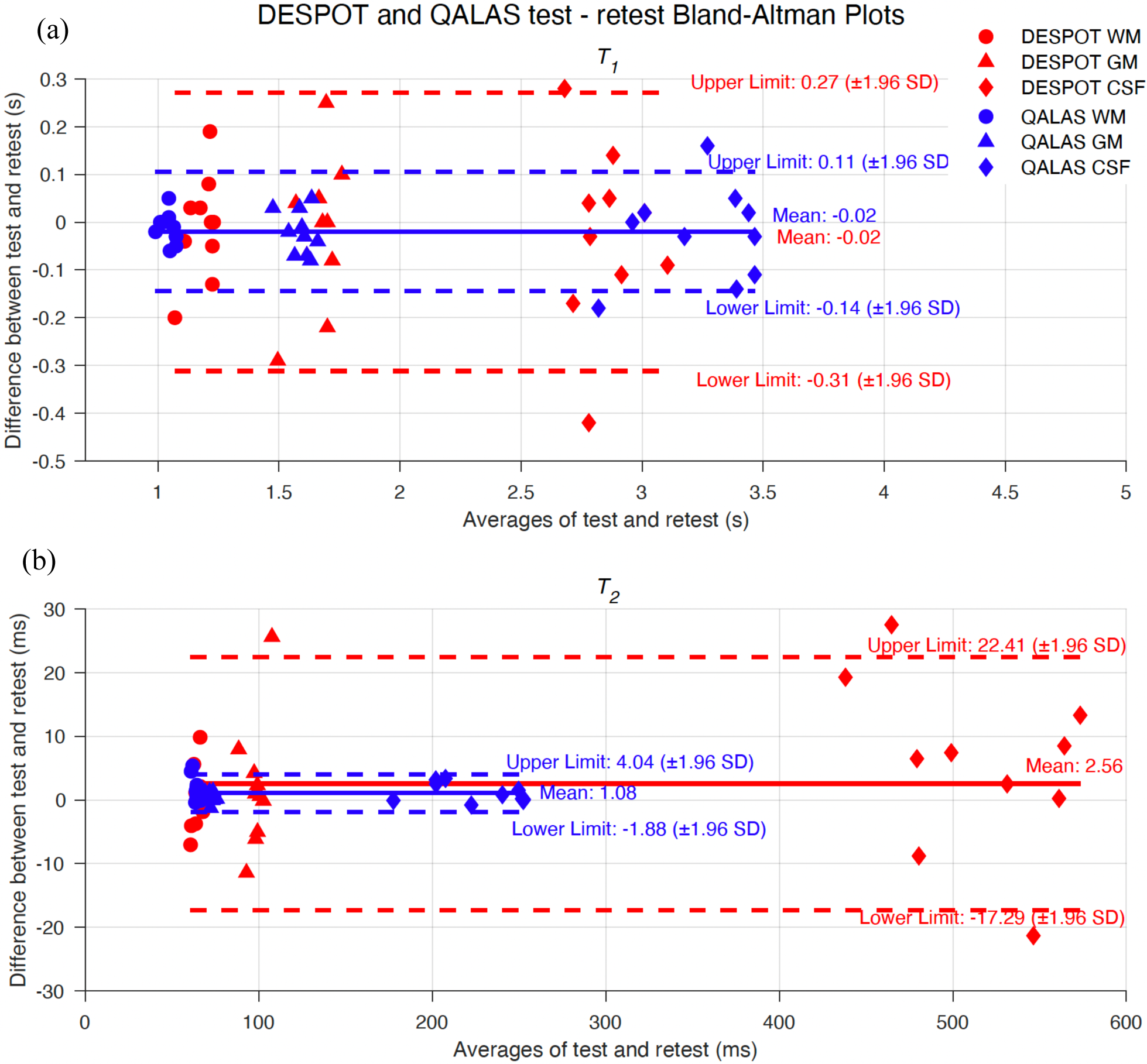
Test-Retest Bland-Altman plots for 10 subjects’ GM, WM and CSF T_1_ and T_2_ values measured by DESPOT and QALAS (a) T_1_ and (b) T_2_. The central solid lines indicate the mean differences (MD), while the upper and lower dotted lines denote the limits of agreement (Mean Difference ± 1.96 * standard deviation (SD) of the differences between test and retest).

#### 3.4. Test-Retest voxel-to-voxel comparisons

Single representative subject test-retest *T*_1_ and *T*_2_ scatter plots for DESPOT and QALAS are shown in Figure 5. The ICCs for all subjects are given in Tables 2 and 3. Higher average voxel-to-voxel ICCs were observed in DESPOT for *T*_2_ (0.901 ± 0.028) and in QALAS for *T*_1_ (0.841 ± 0.039).

**Figure 5:**
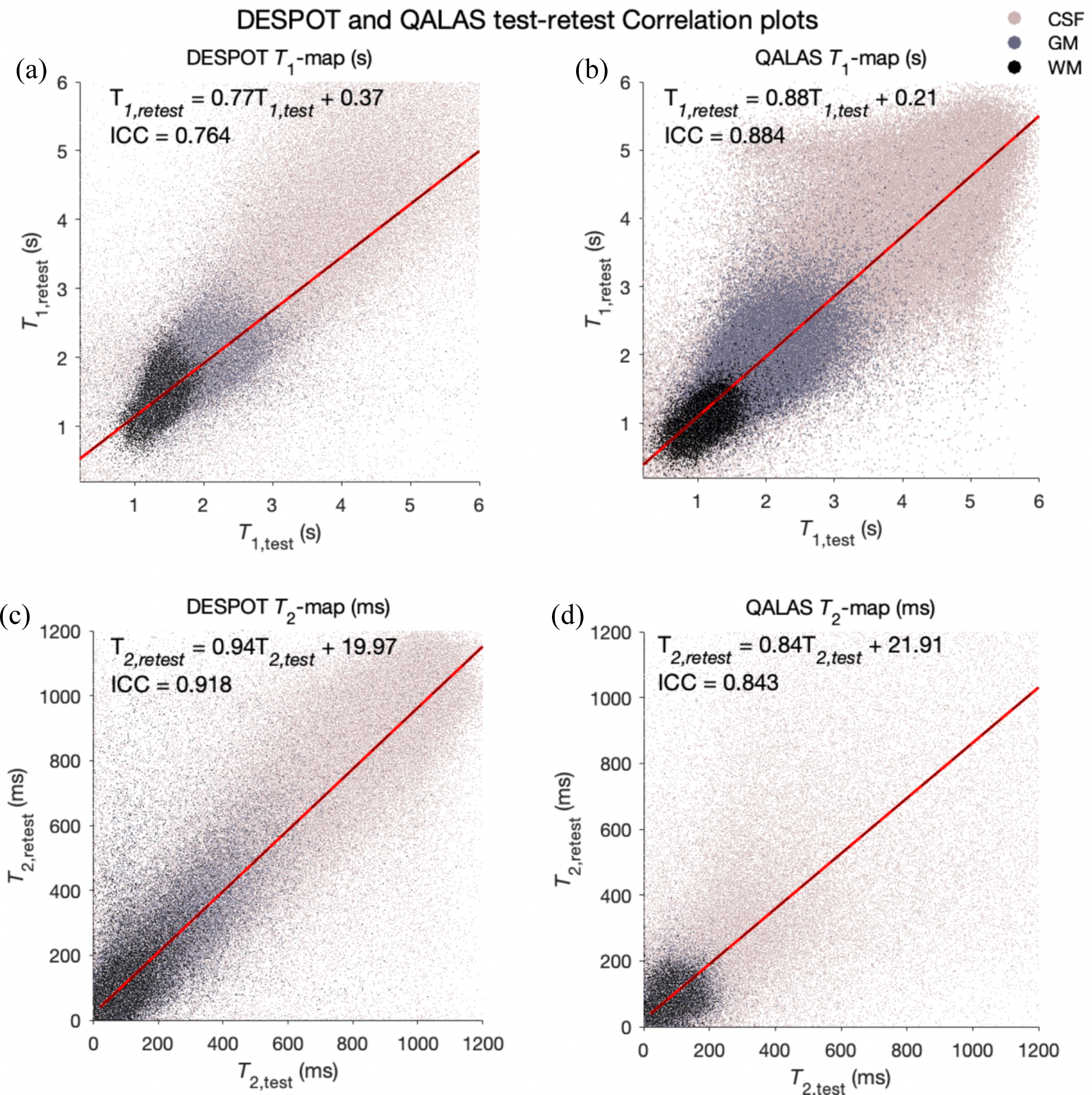
Single-subject test vs. retest scatter plots of T_1_ maps and T_2_ maps. (a) DESPOT T_1_, (b) QALAS T_1_, (c) DESPOT T_2_, and (d) QALAS T_2,_ colored by tissue type. The red diagonal line represents the least-squares regression line.

**Table 2:** DESPOT and QALAS test-retest T_1_ map Voxel to Voxel ICCs.

| Subject | GM |  | WM |  | CSF |  | Whole Brain |  |
| --- | --- | --- | --- | --- | --- | --- | --- | --- |
|  | D | Q | D | Q | D | Q | D | Q |
| 1 | 0.635 | 0.559 | 0.623 | 0.426 | 0.633 | 0.677 | 0.754 | 0.846 |
| 2 | 0.649 | 0.39 | 0.714 | 0.265 | 0.655 | 0.689 | 0.646 | 0.747 |
| 3 | 0.781 | 0.539 | 0.855 | 0.617 | 0.747 | 0.663 | 0.815 | 0.828 |
| 4 | 0.538 | 0.476 | 0.498 | 0.457 | 0.577 | 0.642 | 0.707 | 0.882 |
| 5 | 0.745 | 0.631 | 0.827 | 0.604 | 0.698 | 0.706 | 0.706 | 0.859 |
| 6 | 0.689 | 0.58 | 0.824 | 0.601 | 0.684 | 0.632 | 0.767 | 0.842 |
| 7 | 0.635 | 0.588 | 0.77 | 0.572 | 0.696 | 0.766 | 0.756 | 0.862 |
| 8 | 0.412 | 0.542 | 0.283 | 0.484 | 0.58 | 0.721 | 0.642 | 0.825 |
| 9 | 0.665 | 0.576 | 0.72 | 0.513 | 0.673 | 0.667 | 0.718 | 0.837 |
| 10 | 0.711 | 0.693 | 0.74 | 0.627 | 0.756 | 0.804 | 0.764 | 0.884 |
| Avg ± Std | 0.646 ± 0.106 | 0.557 ± 0.082 | 0.685 ± 0.177 | 0.517 ± 0.114 | 0.670 ± 0.061 | 0.697 ± 0.054 | 0.728 ± 0.055 | 0.841 ± 0.039 |

**Table 3:** DESPOT and QALAS test-retest T_2_ map Voxel to Voxel ICCs.

| Subject | GM |  | WM |  | CSF |  | Whole Brain |  |
| --- | --- | --- | --- | --- | --- | --- | --- | --- |
|  | D | Q | D | Q | D | Q | D | Q |
| 1 | 0.786 | 0.485 | 0.707 | 0.427 | 0.877 | 0.798 | 0.92 | 0.844 |
| 2 | 0.678 | 0.485 | 0.4323 | 0.216 | 0.841 | 0.8 | 0.85 | 0.72 |
| 3 | 0.807 | 0.458 | 0.582 | 0.362 | 0.815 | 0.797 | 0.929 | 0.839 |
| 4 | 0.605 | 0.445 | 0.414 | 0.352 | 0.774 | 0.7462 | 0.852 | 0.7808 |
| 5 | 0.808 | 0.496 | 0.408 | 0.562 | 0.832 | 0.808 | 0.903 | 0.846 |
| 6 | 0.806 | 0.458 | 0.599 | 0.491 | 0.829 | 0.802 | 0.916 | 0.842 |
| 7 | 0.783 | 0.511 | 0.68 | 0.578 | 0.817 | 0.824 | 0.911 | 0.845 |
| 8 | 0.75 | 0.448 | 0.625 | 0.474 | 0.823 | 0.806 | 0.899 | 0.819 |
| 9 | 0.876 | 0.486 | 0.626 | 0.485 | 0.844 | 0.806 | 0.911 | 0.84 |
| 10 | 0.854 | 0.536 | 0.648 | 0.55 | 0.812 | 0.837 | 0.918 | 0.843 |
| Avg ± Std | 0.775 ± 0.081 | 0.481 ± 0.029 | 0.572 ± 0.112 | 0.450 ± 0.113 | 0.826 ± 0.026 | 0.802 ± 0.023 | 0.901 ± 0.028 | 0.822 ± 0.041 |

#### 3.5. Evaluation of agreement between DESPOT and QALAS

Single-subject single-session *T*_1_ and *T*_2_ maps from DESPOT and QALAS and the difference between them are shown in Figure 6. For the same subject and session, the voxel-by-voxel *T*_1_ and *T*_2_ correlation plots between DESPOT and QALAS are shown in Figure 7, suggesting better agreement (ICC) in estimating *T*_1_ values than *T*_2_ values. The Bland-Altman plot for DESPOT against QALAS is depicted in Figure 8. The *T*_1_ plot shows a mean difference of -0.06 s with -0.64 to 0.52 *s* (95% CI) limits of agreement, while the *T*_2_ plot shows a mean difference of 105.29 ms with limits of -157.64 to 368.22 ms, indicating larger variability between the two methods in *T*_2_ measurements compared to *T*_1_, relative to quantity being measured.

**Figure 6:**
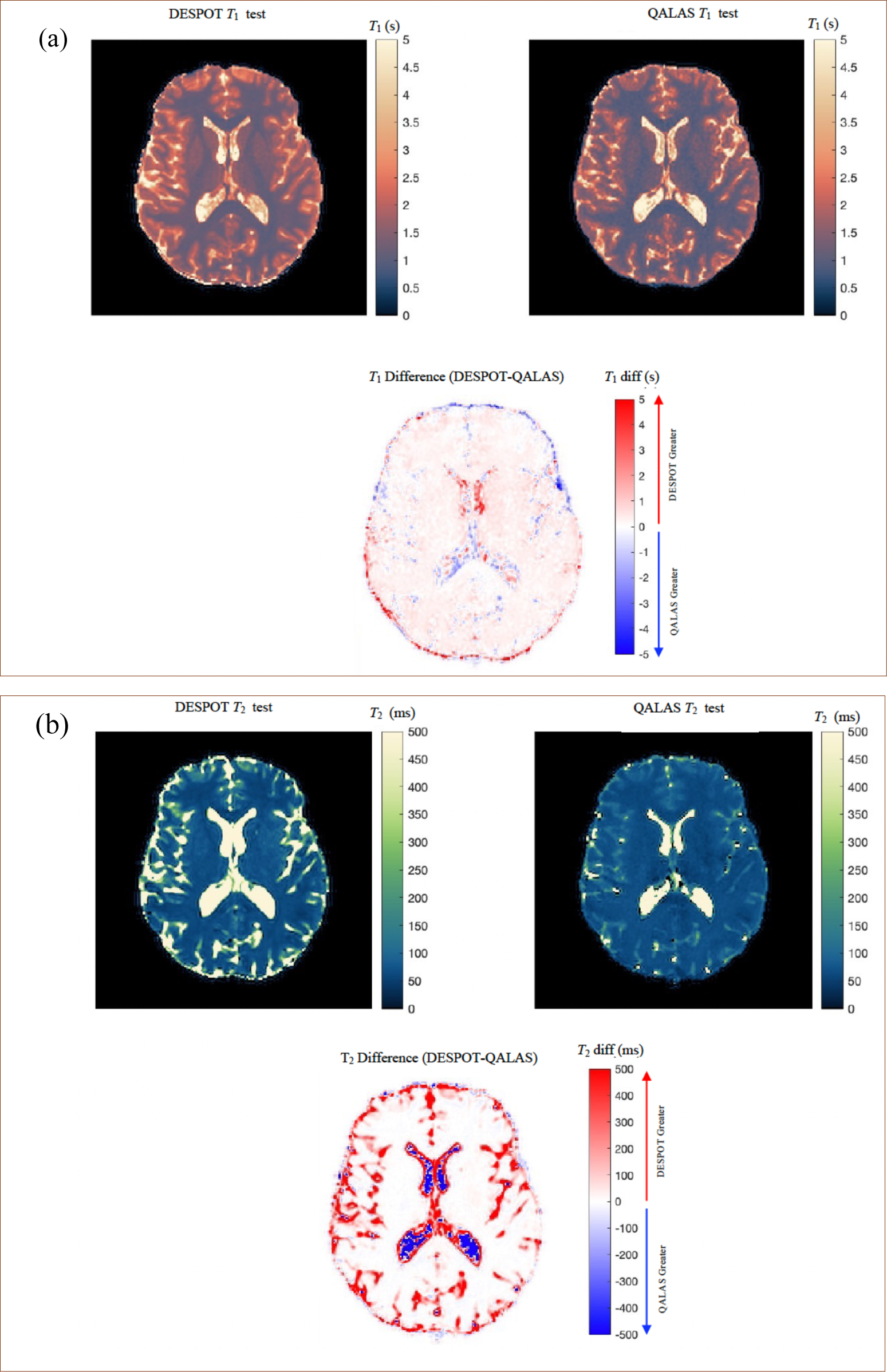
DESPOT and QALAS sample single subject and single session (a) T_1_ and (b) T_2_-maps and difference images.

**Figure 7:**
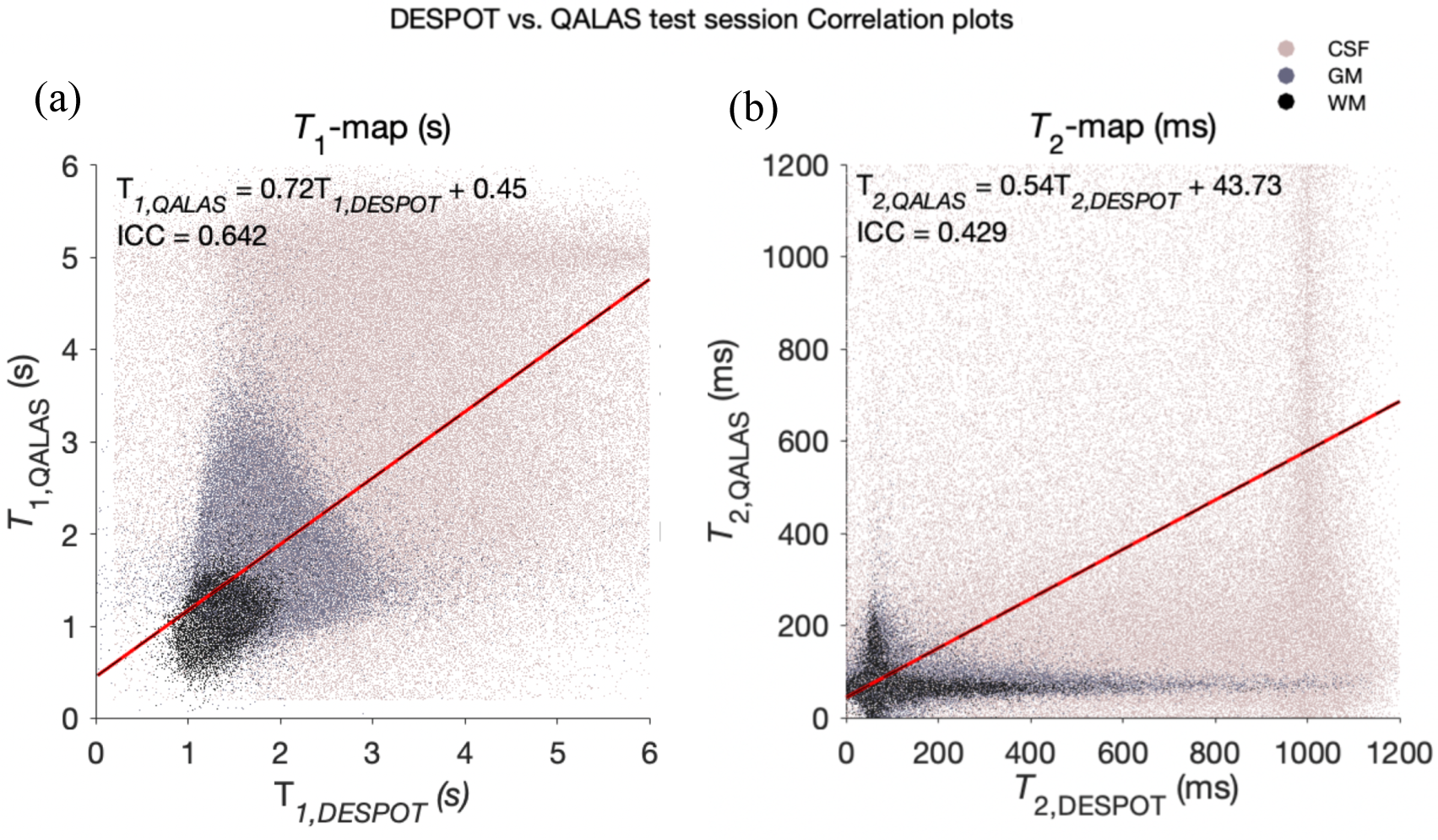
Single subject and single session voxel-to-voxel Test vs Retest scatter plots of DESPOT and QALAS regional T_1_ (left) and T_2_ maps (right). The red line represents the least-squares regression line.

**Figure 8:**
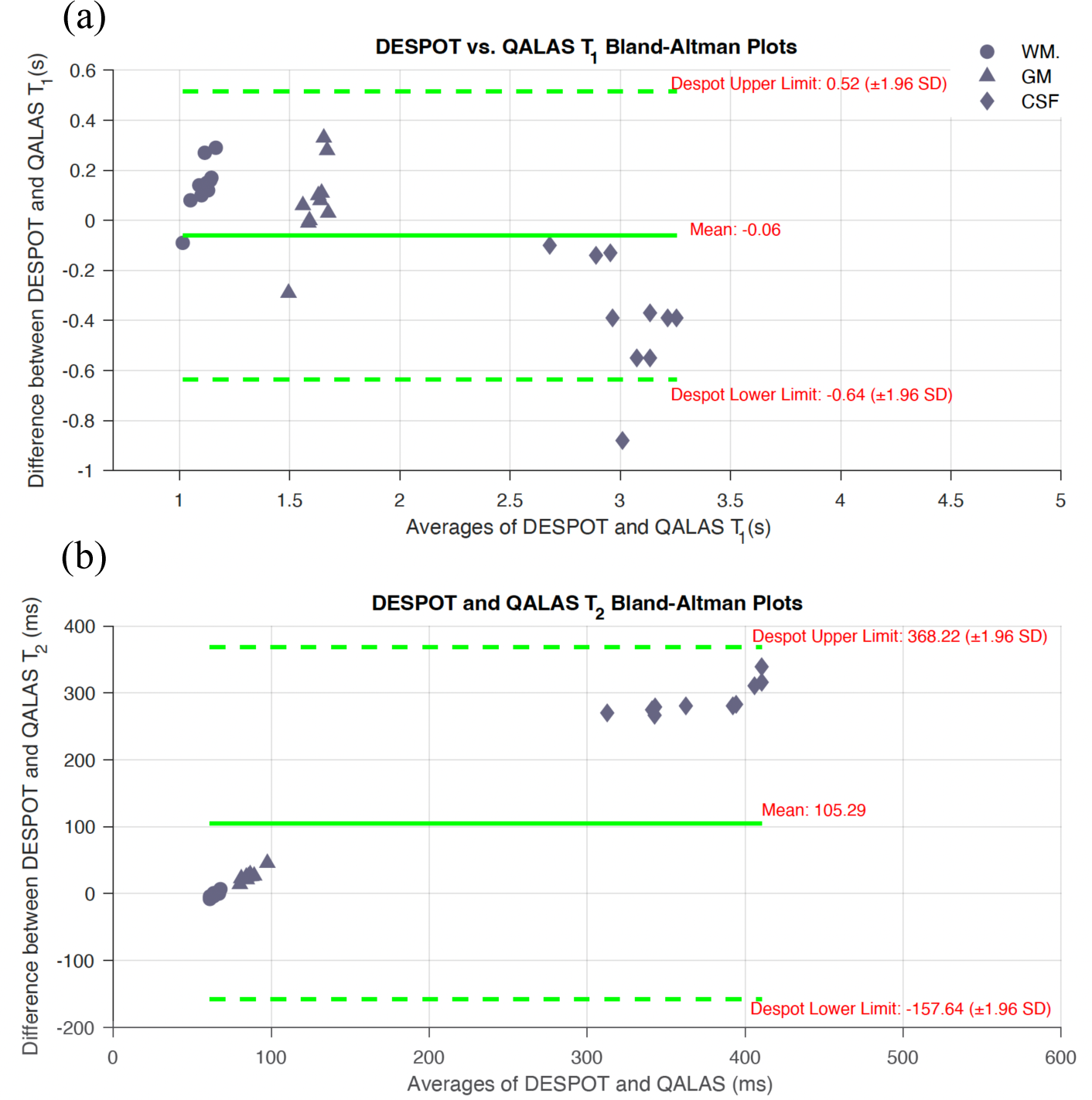
DESPOT vs QALAS Bland-Altman plots for regional mean (a) T_1_ and (b) T_2_ values.

## 4. Discussion

Mapping of *T_1_* and *T_2_* is an invaluable technique in the field of quantitative MR imaging. With a wealth of mapping techniques proposed in the literature, accurately characterizing and comparing the performance of these different methods is paramount to the success of the field. This study specifically investigates the reproducibility of the DESPOT and QALAS techniques for *T_1_* and *T_2_* mapping. Both methods showed good test-retest reproducibility across different brain regions, with some variations in their performance. For *T*_1_ mapping, QALAS exhibited better consistency, both within-subject and between-subject, than DESPOT. The voxel-to-voxel ICCs for *T*_1_ were higher in QALAS, indicating more reliable performance in capturing *T*_1_ values. The Bland-Altman plots for QALAS *T*_1_ and *T*_2_ measurements also showed lower bias and variability than DESPOT. These findings suggest that QALAS may be more suitable for applications where accurate and consistent *T*_1_ mapping is critical, such as in longitudinal studies and clinical assessments of tissue characterization.

It is notable that QALAS achieved slightly better reproducibility than DESPOT, in addition to significantly shorter acquisition time. To generate *T*_1_ and *T*_2_ maps at 1.3 mm^3^ resolution, the current DESPOT protocol acquired 9 separate images. SPGR images at three flip angles are used to determine *T*_1_, while 6 bSSFP images at varying flip angles and phase cycling patterns were used to estimate *T*_2_; acquisition of 0° and 180° phase cycling patterns was performed to address banding artifacts caused by main magnetic field inhomogeneities. While *T*_1_ and *T*_2_ mapping via DESPOT is possible with two varying flip angle SPGR images and two varying flip angle bSSFP images (with phase cycling patterns for *B*_0_-inhomogeneity correction (39)), offering opportunities to make the acquisition more efficient, the accuracy in the *T*_1_ and *T*_2_ model estimates is improved with the additional image information. In contrast, QALAS acquires only 5 images and determines *T*_1_ and *T*_2_ in a single dictionary-matching step. Therefore, QALAS may provide a more efficient acquisition process, using all 5 images to generate both *T*_1_ and *T*_2_ maps, optimizing the utility of acquired data. Furthermore, this single-step process is likely more efficient from the perspective of noise propagation.

The clearest difference between DESPOT and QALAS lies in the *T*_2_ imaging of CSF and partial-volumed CSF areas. Histograms of areas with CSF probability over 0.8 show a large bias – with modal values of ∼1 s for DESPOT and ∼150 ms for QALAS, the longer values being closer to literature precedent (48). Note that the QALAS dictionary does include long *T*_2_s, but for some reason, matching returns much shorter values. DESPOT-QALAS images show large differences in regions for CSF partial voluming, with “partial CSF” returning values resembling CSF in DESPOT, whereas they appear more like GM in QALAS images. The limited resolution of this dataset (1.3 mm^3^) impacts the complete separation of GM from both CSF and WM.

Relaxometry has a number of promising applications in the diagnosis and characterization of several diseases e.g., multiple sclerosis (49), stroke (50), and epilepsy (51,52). Historically, acquisition time was the main barrier to the clinical application of relaxometry mapping techniques, but the advent of rapid acquisition approaches, like those studied here, has circumvented this. The remaining barriers are the accessibility of new techniques, somewhat assuaged by the development of open-source sequence frameworks (53–55), and the establishment of measurement uncertainties, which is the focus of our own study.

The clinical utility of DESPOT for rapid *T_1_* and *T_2_* mapping, emphasizing its speed and feasibility for clinical applications, has been previously studied (33,34). DESPOT’s acquisition scheme makes it attractive for clinical use, especially in scenarios requiring rapid and high-resolution mapping of relaxation times. On the other hand, the capability of QALAS for simultaneous *T_1_* and *T_2_* mapping in the heart (35) and brain (56) has highlighted its efficiency in capturing multiple parameters in a single acquisition. This versatility is extended to GM, WM and CSF brain regions in our study, demonstrating QALAS’s reliability for *T_1_* mapping with good ICC values and minimal bias observed in Bland-Altman plots. This method’s ability to integrate *T_1_*, *T_2_*, and proton density mapping in one short scan makes it a powerful, clinically viable tool for comprehensive tissue characterization.

Our findings add to the body of *T*_1_ and *T*_2_ mapping literature by directly comparing DESPOT and QALAS, demonstrating their respective strengths in reproducibility and mapping accuracy. The DESPOT *T*_1_ values of this study were within acceptable agreement with prior 3T DESPOT *T*_1_ measurements (57) in GM (∼1.6 s) and WM (∼1 s). The DESPOT *T*_2_ values were on average higher than those found in (39) for GM (∼75 ms) and WM (∼50 ms). QALAS *T*_1_ values are significantly longer than in prior 3T QALAS work (58) (∼1.5 vs. ∼1.0 s in GM and ∼1.0 vs ∼0.6 s in WM). QALAS *T*_2_ values are closer to (58) (∼75 vs. 90 ms in GM and ∼65 vs 80 ms in WM). Both our *T*_1_ and *T*_2_ values agree with broader consensus (59), and prior work (44) using a similar analysis pipeline. To the best of our knowledge, there are no reported test-retest studies directly assessing the within-subject reproducibility of DESPOT. However, reproducibility has discussed in terms of percentage standard deviation (SD) of *T*_1_ and *T*_2_ measurements from WM regions (i.e. 6.5 % and 5.5 %) (33) similar to these current findings. One prior QALAS repeatability study (58) showed a mean intrasubject CoV of 1.9 %, similar or slightly better than these findings, although calculated across specific anatomical regions. When compared with other multiparametric mapping techniques, such as ME-MP2RAGE (31,60) and Magnetic Resonance Fingerprinting (MRF) (61,62), the presented methods demonstrated similar within-subject reproducibility. Another multiparameter mapping protocol based on vendor product sequences (i.e. FLASH with MT, T1, and PD contrast weightings) reported an average intra-site CoV of 7 % for *T*_1_ and 16 % for *T*_2_ (63). Taken together, these findings imply both QALAS and DESPOT can be considered robust multiparameter mapping methods.

This is a study of modest scope, with several limitations. Firstly, the study was conducted in a small sample size of 10 healthy volunteers, which may not fully capture the variability in a larger, more diverse population. Secondly, the resolution of mapping was limited to 1.3 mm^3^ isotropic resolution, leading to meaningful partial voluming of cortical GM. Thirdly, only two relaxometry methods are compared. Future studies should include a larger cohort and investigate the reproducibility of these techniques in pathological conditions. Additionally, improvements in the reconstruction algorithms could further enhance the reproducibility and accuracy of both DESPOT and QALAS.

## 5. Conclusion

In summary, this study quantifies the test-retest reproducibility of DESPOT and QALAS for *T*_1_ and *T*_2_ mapping. Given the shorter acquisition time and the slightly better results, QALAS appears to be more reliable, particularly for *T*_1_ measurements, than DESPOT.

## Data and Code Availability

The data used for this study can be available from the corresponding author upon reasonable request

## Author Contributions

GS processed all data, conducted all statistical analyses, prepared all figures, and prepared the manuscript. BG contributed in QALAS protocol development and data processing. YS contributed setup the scan protocol, oversaw the data quality control and data collection procedure. DD contributed to the DESPOT protocol development and data processing. KH and JW contributed to data acquisition protocol and reviewing the manuscript. SM, CDJ, and DS contributed to interpretation of results and drafted parts of the Discussion. VY reviewed all structural scans to assess data quality and check for incidental findings. AG contributed to DESPOT mapping and reviewing the manuscript. HZ and GO contributed to DESPOT and QALAS image analysis and interpretation. RAE designed the project and led interpretation of the results. All authors participated in revision of the manuscript.

## References

1. Ferreira VM, Piechnik SK, Robson MD, Neubauer S, Karamitsos TD. Myocardial Tissue Characterization by Magnetic Resonance Imaging: Novel Applications of T1 and T2 Mapping. J Thorac Imaging. 2014 May;29(3):147.

2. Bohnen S, Radunski UK, Lund GK, Ojeda F, Looft Y, Senel M, et al. Tissue characterization by T1 and T2 mapping cardiovascular magnetic resonance imaging to monitor myocardial inflammation in healing myocarditis. Eur Heart J - Cardiovasc Imaging. 2017 Jul 1;18(7):744–51.

3. Kim PK, Hong YJ, Im DJ, Suh YJ, Park CH, Kim JY, et al. Myocardial T1 and T2 Mapping: Techniques and Clinical Applications. Korean J Radiol. 2017;18(1):113–31.

4. O’Brien AT, Gil KE, Varghese J, Simonetti OP, Zareba KM. T2 mapping in myocardial disease: a comprehensive review. J Cardiovasc Magn Reson. 2022 Jun 6;24(1):33.

5. Mesropyan N, Kupczyk PA, Dold L, Praktiknjo M, Chang J, Isaak A, et al. Assessment of liver cirrhosis severity with extracellular volume fraction MRI. Sci Rep. 2022 Jun 8;12(1):9422.

6. Hoffman DH, Ayoola A, Nickel D, Han F, Chandarana H, Shanbhogue KP. T1 mapping, T2 mapping and MR elastography of the liver for detection and staging of liver fibrosis. Abdom Radiol N Y. 2020 Mar;45(3):692–700.

7. Haimerl M, Verloh N, Zeman F, Fellner C, Müller-Wille R, Schreyer AG, et al. Assessment of Clinical Signs of Liver Cirrhosis Using T1 Mapping on Gd-EOB-DTPA-Enhanced 3T MRI. PLoS ONE. 2013 Dec 31;8(12):e85658.

8. Luetkens JA, Klein S, Träber F, Schmeel FC, Sprinkart AM, Kuetting DLR, et al. Quantification of Liver Fibrosis at T1 and T2 Mapping with Extracellular Volume Fraction MRI: Preclinical Results. Radiology. 2018 Sep;288(3):748–54.

9. Breit HC, Block KT, Winkel DJ, Gehweiler JE, Henkel MJ, Weikert T, et al. Evaluation of liver fibrosis and cirrhosis on the basis of quantitative T1 mapping: Are acute inflammation, age and liver volume confounding factors? Eur J Radiol. 2021 Aug 1;141:109789.

10. Cao G, Gao S, Xiong B. Application of quantitative T1, T2 and T2* mapping magnetic resonance imaging in cartilage degeneration of the shoulder joint. Sci Rep. 2023 Mar 20;13(1):4558.

11. Mittal S, Pradhan G, Singh S, Batra R. T1 and T2 mapping of articular cartilage and menisci in early osteoarthritis of the knee using 3-Tesla magnetic resonance imaging. Pol J Radiol. 2019;84(1):549–64.

12. Zhao H, Li H, Liang S, Wang X, Yang F. T2 mapping for knee cartilage degeneration in young patients with mild symptoms. BMC Med Imaging. 2022 Apr 18;22(1):72.

13. Shah NJ, Neeb H, Zaitsev M, Steinhoff S, Kircheis G, Amunts K, et al. Quantitative T1 mapping of hepatic encephalopathy using magnetic resonance imaging. Hepatology. 2003 Nov;38(5):1219–26.

14. Vrenken H, Geurts JJG, Knol DL, van Dijk LN, Dattola V, Jasperse B, et al. Whole-Brain T1 Mapping in Multiple Sclerosis: Global Changes of Normal-appearing Gray and White Matter. Radiology. 2006 Sep;240(3):811–20.

15. Lescher S, Jurcoane A, Veit A, Bähr O, Deichmann R, Hattingen E. Quantitative T1 and T2 mapping in recurrent glioblastomas under bevacizumab: earlier detection of tumor progression compared to conventional MRI. Neuroradiology. 2015 Jan 1;57(1):11–20.

16. Müller A, Jurcoane A, Kebir S, Ditter P, Schrader F, Herrlinger U, et al. Quantitative T1-mapping detects cloudy-enhancing tumor compartments predicting outcome of patients with glioblastoma. Cancer Med. 2017;6(1):89–99.

17. Eminian S, Hajdu SD, Meuli RA, Maeder P, Hagmann P. Rapid high resolution T1 mapping as a marker of brain development: Normative ranges in key regions of interest. PLOS ONE. 2018 Jun 14;13(6):e0198250.

18. Grossman R, Hoffman C, Mardor Y, Biegon A. Quantitative MRI measurements of human fetal brain development *in utero*. NeuroImage. 2006 Nov 1;33(2):463–70.

19. Giedd JN, Snell JW, Lange N, Rajapakse JC, Casey BJ, Kozuch PL, et al. Quantitative magnetic resonance imaging of human brain development: ages 4-18. Cereb Cortex N Y N 1991. 1996;6(4):551– 60.

20. Kumar R, Delshad S, Woo MA, Macey PM, Harper RM. Age-related regional brain T2-relaxation changes in healthy adults. J Magn Reson Imaging JMRI. 2012 Feb;35(2):300–8.

21. Callaghan MF, Freund P, Draganski B, Anderson E, Cappelletti M, Chowdhury R, et al. Widespread age-related differences in the human brain microstructure revealed by quantitative magnetic resonance imaging. Neurobiol Aging. 2014 Aug 1;35(8):1862–72.

22. Carey D, Caprini F, Allen M, Lutti A, Weiskopf N, Rees G, et al. Quantitative MRI provides markers of intra-, inter-regional, and age-related differences in young adult cortical microstructure. NeuroImage. 2018 Nov 15;182:429–40.

23. Studler U, White LM, Andreisek G, Luu S, Cheng HLM, Sussman MS. Impact of motion on T1 mapping acquired with inversion recovery fast spin echo and rapid spoiled gradient recalled-echo pulse sequences for delayed gadolinium-enhanced MRI of cartilage (dGEMRIC) in volunteers. J Magn Reson Imaging. 2010;32(2):394–8.

24. Messroghli DR, Radjenovic A, Kozerke S, Higgins DM, Sivananthan MU, Ridgway JP. Modified Look-Locker inversion recovery (MOLLI) for high-resolution T1 mapping of the heart. Magn Reson Med. 2004;52(1):141–6.

25. Piechnik SK, Ferreira VM, Dall’Armellina E, Cochlin LE, Greiser A, Neubauer S, et al. Shortened Modified Look-Locker Inversion recovery (ShMOLLI) for clinical myocardial T1-mapping at 1.5 and 3 T within a 9 heartbeat breathhold. J Cardiovasc Magn Reson Off J Soc Cardiovasc Magn Reson. 2010 Nov 19;12(1):69.

26. Chow K, Flewitt JA, Green JD, Pagano JJ, Friedrich MG, Thompson RB. Saturation recovery single-shot acquisition (SASHA) for myocardial T(1) mapping. Magn Reson Med. 2014 Jun;71(6):2082–95.

27. Slavin GS, Stainsby JA. True T1 mapping with SMART1Map (saturation method using adaptive recovery times for cardiac T1 mapping): a comparison with MOLLI. J Cardiovasc Magn Reson. 2013 Jan 30;15(Suppl 1):P3.

28. Schmitt P, Griswold MA, Jakob PM, Kotas M, Gulani V, Flentje M, et al. Inversion recovery TrueFISP: quantification of T(1), T(2), and spin density. Magn Reson Med. 2004 Apr;51(4):661–7.

29. Fram EK, Herfkens RJ, Johnson GA, Glover GH, Karis JP, Shimakawa A, et al. Rapid calculation of T1 using variable flip angle gradient refocused imaging. Magn Reson Imaging. 1987;5(3):201–8.

30. Heule R, Ganter C, Bieri O. Variable flip angle T1 mapping in the human brain with reduced T2 sensitivity using fast radiofrequency-spoiled gradient echo imaging. Magn Reson Med. 2016 Apr;75(4):1413–22.

31. Marques JP, Kober T, Krueger G, van der Zwaag W, Van de Moortele PF, Gruetter R. MP2RAGE, a self bias-field corrected sequence for improved segmentation and T1-mapping at high field. NeuroImage. 2010 Jan 15;49(2):1271–81.

32. Caan MWA, Bazin P, Marques JP, de Hollander G, Dumoulin SO, van der Zwaag W. MP2RAGEME: T1, T2 *, and QSM mapping in one sequence at 7 tesla. Hum Brain Mapp. 2018 Dec 13;40(6):1786– 98.

33. Deoni SCL, Rutt BK, Peters TM. Rapid combined T1 and T2 mapping using gradient recalled acquisition in the steady state. Magn Reson Med. 2003;49(3):515–26.

34. Deoni SCL, Peters TM, Rutt BK. High-resolution T1 and T2 mapping of the brain in a clinically acceptable time with DESPOT1 and DESPOT2. Magn Reson Med. 2005;53(1):237–41.

35. Kvernby S, Warntjes MJB, Haraldsson H, Carlhäll CJ, Engvall J, Ebbers T. Simultaneous three-dimensional myocardial T1 and T2 mapping in one breath hold with 3D-QALAS. J Cardiovasc Magn Reson. 2014 Dec 20;16(1):102.

36. Roujol S, Weingärtner S, Foppa M, Chow K, Kawaji K, Ngo LH, et al. Accuracy, Precision, and Reproducibility of Four T1 Mapping Sequences: A Head-to-Head Comparison of MOLLI, ShMOLLI, SASHA, and SAPPHIRE. Radiology. 2014 Sep;272(3):683–9.

37. Moon JC, Messroghli DR, Kellman P, Piechnik SK, Robson MD, Ugander M, et al. Myocardial T1 mapping and extracellular volume quantification: a Society for Cardiovascular Magnetic Resonance (SCMR) and CMR Working Group of the European Society of Cardiology consensus statement. J Cardiovasc Magn Reson. 2013 Oct 14;15(1):92.

38. Kellman P, Hansen MS. T1-mapping in the heart: accuracy and precision. J Cardiovasc Magn Reson. 2014 Jan 4;16(1):2.

39. Deoni SCL. Transverse relaxation time (T2) mapping in the brain with off-resonance correction using phase-cycled steady-state free precession imaging. J Magn Reson Imaging. 2009;30(2):411–7.

40. Yarnykh VL. Actual flip-angle imaging in the pulsed steady state: A method for rapid three-dimensional mapping of the transmitted radiofrequency field. Magn Reson Med. 2007;57(1):192–200.

41. Jordan CJ, Weiss SRB, Howlett KD, Freund MP. Introduction to the Special Issue on “Informing Longitudinal Studies on the Effects of Maternal Stress and Substance Use on Child Development: Planning for the HEALthy Brain and Child Development (HBCD) Study.” Advers Resil Sci. 2020 Dec 1;1(4):217–21.

42. Jenkinson M, Bannister P, Brady M, Smith S. Improved optimization for the robust and accurate linear registration and motion correction of brain images. NeuroImage. 2002 Oct;17(2):825–41.

43. Dean III DC, Adluru N, Guerrero J. QMRI-neuropipe: A flexible software framework for the analysis of quantitative MRI data. In: Proc Intl Soc Mag Reson Med. 2023. p. 2406.

44. Cho J, Gagoski B, Kim TH, Wang F, Manhard MK, Dean III D, et al. Time-efficient, high-resolution 3T whole-brain relaxometry using 3D-QALAS with wave-CAIPI readouts. Magn Reson Med. 2024;91(2):630–9.

45. Ben-Eliezer N, Sodickson DK, Block KT. Rapid and accurate T2 mapping from multi–spin-echo data using Bloch-simulation-based reconstruction. Magn Reson Med. 2015;73(2):809–17.

46. Ma D, Coppo S, Chen Y, McGivney DF, Jiang Y, Pahwa S, et al. Slice profile and B1 corrections in 2D magnetic resonance fingerprinting. Magn Reson Med. 2017 Nov;78(5):1781–9.

47. Ashburner J, Friston KJ. Unified segmentation. NeuroImage. 2005 Jul 1;26(3):839–51.

48. Spijkerman JM, Petersen ET, Hendrikse J, Luijten P, Zwanenburg JJM. T2 mapping of cerebrospinal fluid: 3 T versus 7 T. Magma N Y N. 2018;31(3):415–24.

49. Granziera C, Wuerfel J, Barkhof F, Calabrese M, De Stefano N, Enzinger C, et al. Quantitative magnetic resonance imaging towards clinical application in multiple sclerosis. Brain. 2021 May 1;144(5):1296– 311.

50. Bernarding J, Braun J, Hohmann J, Mansmann U, Hoehn-Berlage M, Stapf C, et al. Histogram-based characterization of healthy and ischemic brain tissues using multiparametric MR imaging including apparent diffusion coefficient maps and relaxometry. Magn Reson Med. 2000;43(1):52–61.

51. Chen H, Yu G, Wang J, Li F, Li G. Application of T2 relaxometry in lateralization and localization of mesial temporal lobe epilepsy and corresponding comparison with MR volumetry. Acta Radiol. 2016 Sep 1;57(9):1107–13.

52. Pell GS, Briellmann RS, Waites AB, Abbott DF, Jackson GD. Voxel-based relaxometry: a new approach for analysis of T2 relaxometry changes in epilepsy. NeuroImage. 2004 Feb 1;21(2):707–13.

53. Karakuzu A, Biswas L, Cohen-Adad J, Stikov N. Vendor-neutral sequences and fully transparent workflows improve inter-vendor reproducibility of quantitative MRI. Magn Reson Med. 2022;88(3):1212–28.

54. Layton KJ, Kroboth S, Jia F, Littin S, Yu H, Leupold J, et al. Pulseq: A rapid and hardware-independent pulse sequence prototyping framework. Magn Reson Med. 2017;77(4):1544–52.

55. Cordes C, Konstandin S, Porter D, Günther M. Portable and platform-independent MR pulse sequence programs. Magn Reson Med. 2020;83(4):1277–90.

56. Yamashita K, Yoneyama M, Kikuchi K, Wada T, Murazaki H, Watanuki H, et al. Reproducibility of quantitative ADC, T1, and T2 measurement on the cerebral cortex: Utility of whole brain echo-planar DWI with compressed SENSE (EPICS-DWI): A pilot study. Eur J Radiol Open. 2023 Dec 1;11:100516.

57. Deoni SCL. High-resolution T1 mapping of the brain at 3T with driven equilibrium single pulse observation of T1 with high-speed incorporation of RF field inhomogeneities (DESPOT1-HIFI). J Magn Reson Imaging. 2007;26(4):1106–11.

58. Fujita S, Hagiwara A, Hori M, Warntjes M, Kamagata K, Fukunaga I, et al. Three-dimensional high-resolution simultaneous quantitative mapping of the whole brain with 3D-QALAS: An accuracy and repeatability study. Magn Reson Imaging. 2019 Nov 1;63:235–43.

59. Stikov N, Boudreau M, Levesque IR, Tardif CL, Barral JK, Pike GB. On the accuracy of T1 mapping: searching for common ground. Magn Reson Med. 2015 Feb;73(2):514–22. doi: 10.1002/mrm.25135. Epub 2014 Feb 27. PMID: 24578189.

60. Metere R, Kober T, Möller HE, Schäfer A. Simultaneous Quantitative MRI Mapping of T1, T2* and Magnetic Susceptibility with Multi-Echo MP2RAGE. PLOS ONE. 2017 Jan 12;12(1):e0169265.

61. Buonincontri G, Biagi L, Retico A, Cecchi P, Cosottini M, Gallagher FA, et al. Multi-site repeatability and reproducibility of MR fingerprinting of the healthy brain at 1.5 and 3.0 T. NeuroImage. 2019 Jul 15;195:362–72.

62. Körzdörfer G, Kirsch R, Liu K, Pfeuffer J, Hensel B, Jiang Y, et al. Reproducibility and Repeatability of MR Fingerprinting Relaxometry in the Human Brain. Radiology. 2019 Aug;292(2):429– 37.

63. Leutritz T, Seif M, Helms G, Samson RS, Curt A, Freund P, et al. Multiparameter mapping of relaxation (R1, R2*), proton density and magnetization transfer saturation at 3 T: A multicenter dual-vendor reproducibility and repeatability study. Hum Brain Mapp. 2020;41(15):4232–47.

